# Selective injury of thalamocortical tract in neonatal rats impairs forelimb use: model validation and behavioral effects

**DOI:** 10.1101/2024.08.07.607003

**Authors:** Tong Chun Wen, Michelle Corkrum, Jason B. Carmel

**Affiliations:** Movement Recovery Laboratory, Department of Neurology, Columbia University Medical Center, New York, New York, USA

**Keywords:** Sensory, motor, injury, movement, neonatal

## Abstract

Unilateral brain injury in neonates results in largely contralateral hand function in children. Most research investigating neurorehabilitation targets for movement recovery has focused on the effects of brain injury on descending motor systems, especially the corticospinal tract. However, a recent human study demonstrated that sensory tract injury may have larger effects on dexterity than motor tract injury. To validate that the sensory tract injury impairs dexterity, we modeled the most common site of sensory tract injury in neonates by targeting the thalamocortical tract. In the postnatal day 7 rats, we used three types of lesions to the thalamocortical tract: periventricular blood injection, photothrombotic lesion, and electrolytic lesion. To test the sensitivity and specificity of these techniques, viral tracers were injected into the primary sensory or motor cortex immediately after injury. Electrolytic lesions were the most specific and reproducible for inducing a lesion compared to the other two methods. Electrolytic lesions disrupted 63% of the thalamocortical tract, while sparing the adjacent corticospinal tract in the internal capsule. To measure the impact on dexterity, the cylinder exploration and pasta handling tests were used to test the changes of forelimb use at 8 weeks after injury when the rats reached maturity. Lesions to the thalamocortical tract were associated with a significant decrease in the use of the contralateral forelimb in the cylinder task, and the degree of impairment positively correlated with the degree of injury. Overall, specific sensory system lesions of the thalamocortical tract impair forelimb use, suggesting a key role for skilled movement.

## Introduction

Congenital hemiplegia is commonly caused by injury to one hemisphere and primarily leads to impaired hand function on the side contralateral to brain injury. This can occur at pre-term or full-term, and affects1-2/1000 children [1]. The impairment is life long and diminishes participation in activities and quality of life [2, 3]. Therefore, improving hand function is usually the highest priority for people with congenital hemiplegia [4–6].

Skilled movement requires descending motor control and sensory feedback. Although previous research has primarily focused on the descending corticospinal tracts for movement recovery, emerging research has revealed a crucial role of sensory tracts in congenital hemiplegia. It is thought that corticospinal injury has less pronounced effect on dexterity because of the persistence of bilateral connections from the uninjured hemisphere [7, 8] if the injury is severe or early in development [9]. Unilateral brain injury produces substantial sensory deficits in children with congenital hemiplegia [10–12], demonstrating that skilled movement requires descending motor control and sensory feedback.

Previously, we investigated the relative contribution of sensory and motor connectivity for hand function in children with hemiplegia [10]. In this study, two groups of school-aged children were compared, those with subcortical periventricular injury that primarily injured the corticospinal tract (CST) and often spared the thalamocortical sensory tract (TCT), and those with middle cerebral artery (MCA) injuries that often had injury to both motor and sensory cortices and their subcortical tracts. We measured the anatomy and physiology of motor and sensory connections and compared each of these with hand function. For anatomy, tractographies of the CST and the medial lemniscus (ML) were measured with MRI. For physiology, motor evoked responses to transcranial magnetic stimulation and somatosensory evoked potentials to finger vibration were measured. Hand functions were assessed with tests of movement (Assisting Hand Assessment, Jebsen-Taylor, and Box & Blocks) and sensation (2-point discrimination and stereognosis). In the multivariate analysis of these variables (MANOVA), only three variables were independently associated with hand functions: sensory physiology (p=0.0003; Eta2 0.70), sensory anatomy (0.0005; 0.37), and lesion type (0.0005; 0.42). Effect size measured by Eta2 is considered large if it is over 0.26; thus, these effect sizes are considered very large. These data strongly suggest that sensory connectivity most strongly limits hand function. In addition, MCA lesions were more strongly associated with poor hand function than periventricular lesions, as others have also shown [13]. Importantly, researchers have shown that the ascending TCT contributes to hand function in both types of injury [10,14]. However, more research is needed for accurate and efficacious targeting of sensory circuits and sufficient target engagement in terms of manipulating the connectivity and function of sensory circuits. To close these knowledge gaps, it is critical to develop a reliable preclinical stroke model that examines the effect of sensory only injury.

## Methods

In this study, we measured the effect of forelimb preference after selective TCT lesion. To develop such a rodent model, we considered the need to mimic pathophysiology observed in pediatric patients and cause a highly selective lesion to TCT. The location is a clinically important site for brain injury, especially in children born prematurely [15]. It also allows comparison with the adjacent CST in the internal capsule (IC). A challenge for developing this model is that it is difficult to fix neonatal rats on the stereotaxic frame accurately because their skulls have not fully ossified, resulting in error in targeting indicated coordinates. Secondly, the sensory and the motor tracts course closely together in the brain (Figure 1), increasing the difficulty of specifically targeting the sensory tract while sparing the motor tract. The available approaches for inducing this model include chemical injection, blood vessel occlusion, blood injection, photothrombosis, and electrolytic technique. It is well known that chemical injection can diffuse in unpredictable ways including along the injection tract and can cause off-target injury [16]. Blood vessel occlusion such as MCA occlusion affects both the sensory and the motor tracts [17,18]. Therefore, in the current study, we used the three other approaches to induce injury to TCT in P7 rats. First, we tried the blood injection method, which most closely mimics the clinical condition of periventricular hemorrhagic infarction [19]. Then we tested the photothrombotic method, which mimics the ischemic component. Finally, we used the electrolytic method, which can cause highly selective lesions to TCT, but causes injury largely through electrical or thermal injury rather than ischemia. To assess the effects of sensory lesion on paw use, we used two behavioral tasks, the cylinder exploration task and the pasta handling task.

**Figure 1.**
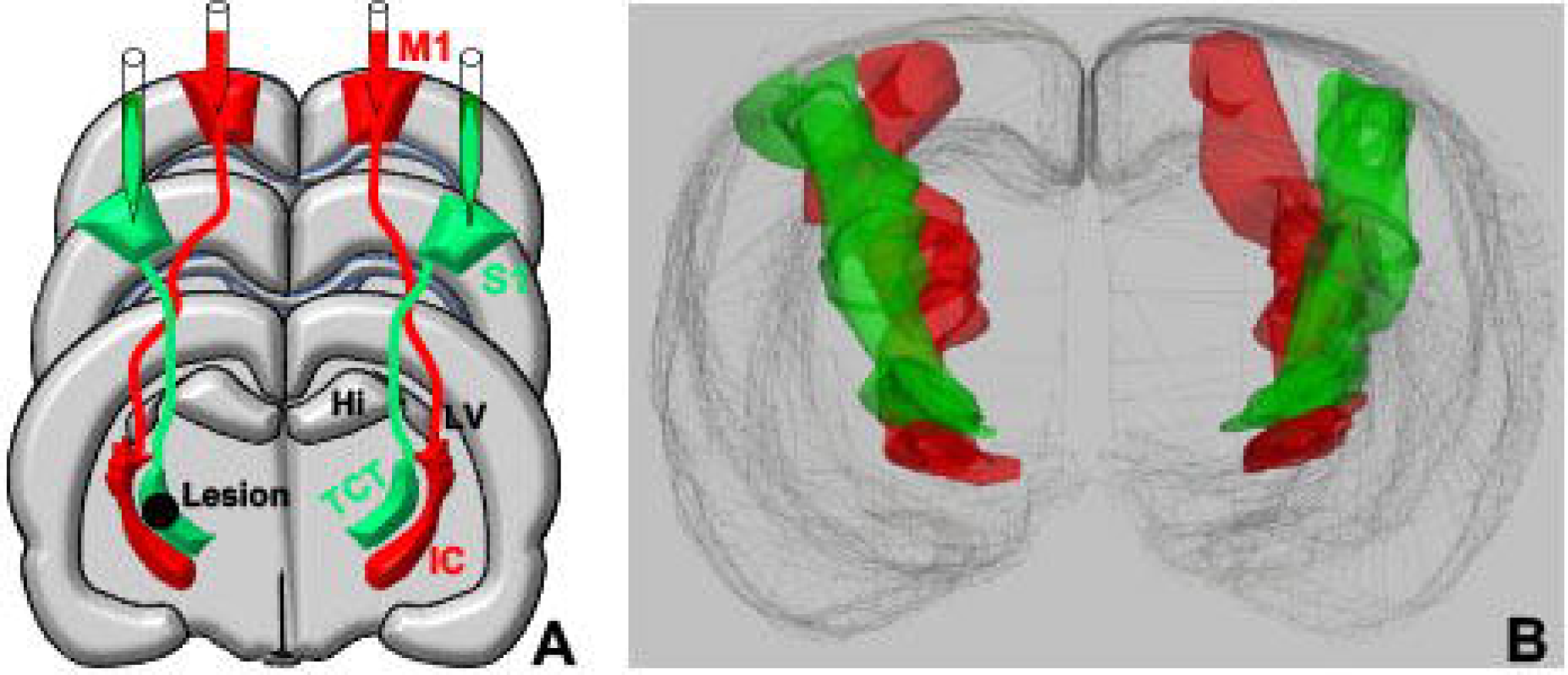
Setup of the experiment. **A**, The targeted lesioned area (thalamocortical tracts, TCT) and tracing. The retrograde adeno-associated virus 2-enhanced green fluorescent protein (AAV2-rGFP) was injected into the sensory cortex (S1) while the anterograde tracer AAV8-CAG.CI-rCOMET (AAV8-rCOMET) was injected into the motor cortex (M1). Hi, Hippocampus, IC, internal capsule, LV, lateral ventricle. **B,** 3D reconstruction the TCT (green, AAV2-rGFP) and CST (red, AAV8-rCOMET).

### Institutional approvals

The study was carried out in accordance with the recommendations of National Institutes of Health guidelines and was approved by the Institutional Animal Care and Use Committee of Columbia University.

### Animals

Postnatal day 5 (P5) Sprague-Dawley rat pups with the dam were purchased from Charles River Laboratories (Wilmington, MA, USA), and housed in an animal care facility with food and water ad libitum. The lights were timed so that the housing room was dark during the day (9 am– 9 pm) to facilitate behavior testing during the nocturnal period. Brain lesions were performed at P7, an age at which the developmental stage of the rat brain is histologically similar to that of the 34–35-week gestation human [20, 21]. A total of 78 rat pups were used in the study, 24 to work out the parameters of the injury models and 54 for the experiments measuring the anatomical and behavioral outcomes. Among 54 rat pups, 20 were used for blood injection, 8 for photothrombotic lesion, 18 for electrolytic lesion (12 active lesion and 6 for sham controls that placed the electrode but did not deliver the electrolytic lesion), and 8 for the behavioral tests after electrolytic lesion. The preliminary experiments were performed to optimize our techniques for electrolytic lesion. We tested the microelectrode size (N=12), and the current intensity and time (N=12). For a microelectrode size, we tested four different diameters of the microelectrode: 125, 250, 500, and 750 µm. For the current intensity and time, based on the studies of Ortiz-Pulido R, et al [22] and Ouhaz Z et al [23], we tested 0.5mA current for 10 or 20 s and 5 mA for 3 or 10 s. After analysis of the lesions (not shown), we chose a microelectrode of 500 µm in diameter and 0.5 mA current for 10 s to cause the most selective lesions.

### Blood injection

The rat pups (N=20) were anesthetized using 1%–1.5% isoflurane in oxygen (Euthanex Corp., Palmer, PA, USA) at a flow rate of 1 liter/min and fixed on a neonatal rat stereotaxic frame (Stoelting Co., Wood Dale, IL, USA). Autologous blood (25 ul) was collected into a sterile syringe from the tail after cleaning the tail skin with 70% alcohol and cutting 2 mm off the tail tip. A small craniotomy was made at 1.2 mm posterior to Bregma and 5.0 mm lateral to midline. A needle attached to the syringe was introduced via the craniotomy under ultrasound guidance (Figure 2A). The lesion target was the periventricular area (Figure 2B). Ultrasound guidance was performed using a VisualSonics Vevo2100 High Resolution Ultrasound (VisualSonics, Toronto, Canada). Non-toxic ultrasound gel was applied to the skin during the procedure. Following blood injection, the wound was closed with topical tissue adhesive, and the pups were warmed on a heating plate until fully awake before being returned to their dam.

**Figure 2.**
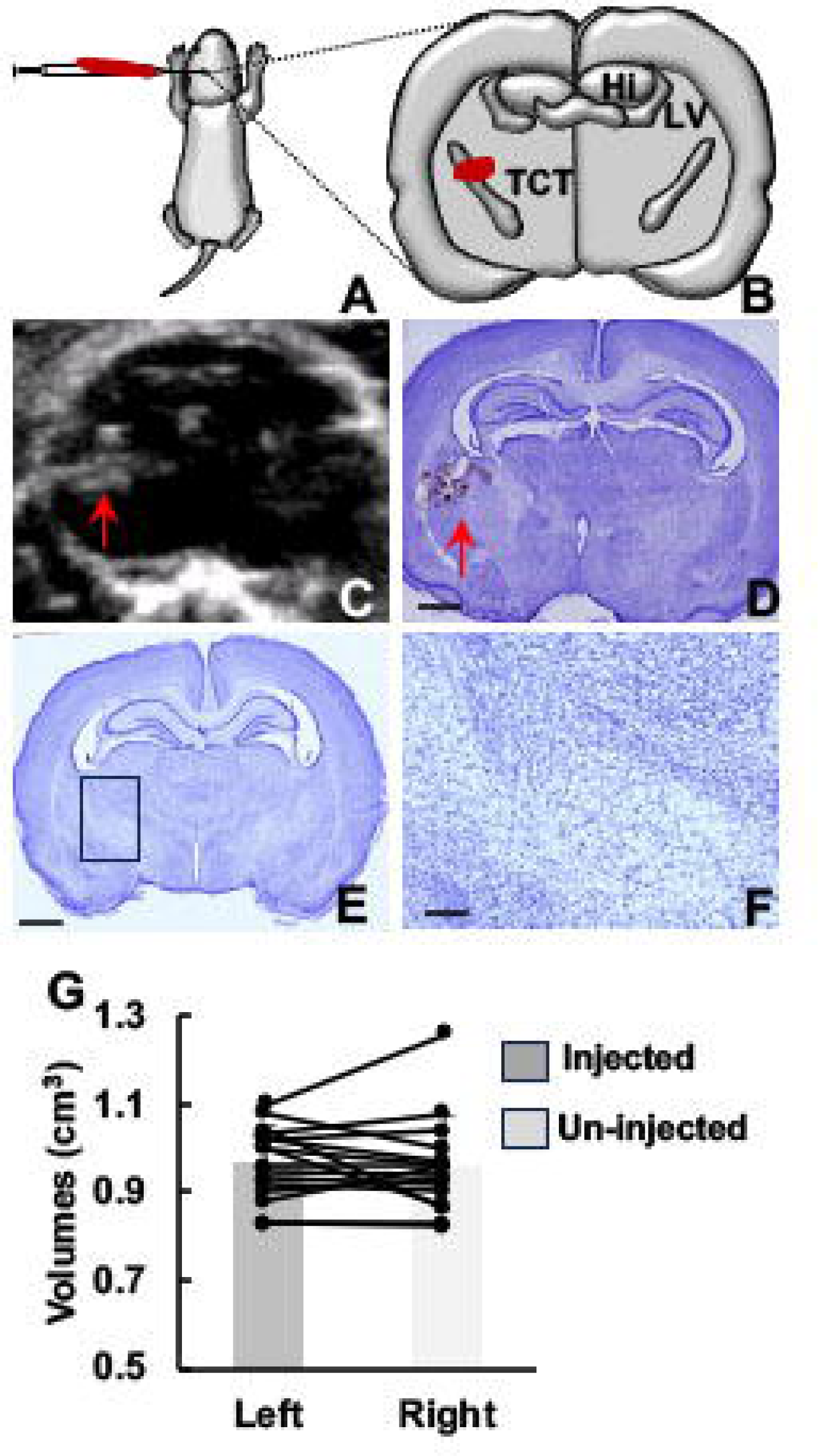
Blood injection. **A,** Schematic of blood injection. **B,** The target area of blood injection. **C,** An ultrasound image of the blood injection (red arrow). **D,** Hematoma at 3 days after blood injection (red arrow). **E,** Nissl staining at 12 weeks after blood injection. **F**, High magnification of the injection side. No injury was detected. **D,** Volumes of each hemisphere in the rats at 12 weeks after blood injection. Bar = 500 μm in D and E, 125 μm in F.

To confirm the size and location of the hematoma after blood injection, the rats (N=4) were perfused transcardially with 0.1 M phosphate buffer saline (PBS), followed by 4% paraformaldehyde in 0.1 M phosphate buffer 3 days after blood injection. For long-term effects of blood injection, rats (N=16) were perfused at 12 weeks after blood injection. After perfusion, rat brains were removed, postfixed overnight in 4% paraformaldehyde, and then transferred to a solution of 30% sucrose in 0.1 M PBS for 3 days. The brain region of the injection was blocked, frozen with dry ice, and kept at −20 ℃ until sectioning. Coronal sections (40 μm) were cut on a cryostat and stained with 0.1% cresyl violet (Abcam, Cambridge, United Kingdom).

### Photothrombotic lesions

P7 rat pups (N=8) were anesthetized with isoflurane and fixed in the neonatal rat stereotaxic frame (Stoelting Co.). A small craniotomy was made at 1.2 mm posterior to Bregma and 3.0 mm lateral to midline. An optical fiber (diameter: 125 um, Doric Lenses, Quebec, Canada) was introduced into the rat brain (3.5 mm deep to the skull surface) via the craniotomy (Figure 3A). Rose Bengal was injected intraperitoneally (20 mg/kg) followed 20 minutes later by delivery of light (the current 0.55 mA, LD Fiber Light Source, Doric Lenses) for 90 seconds. We started by injecting the dye into the vein, but found intraperitoneal injection was more reproducible.

**Figure 3.**
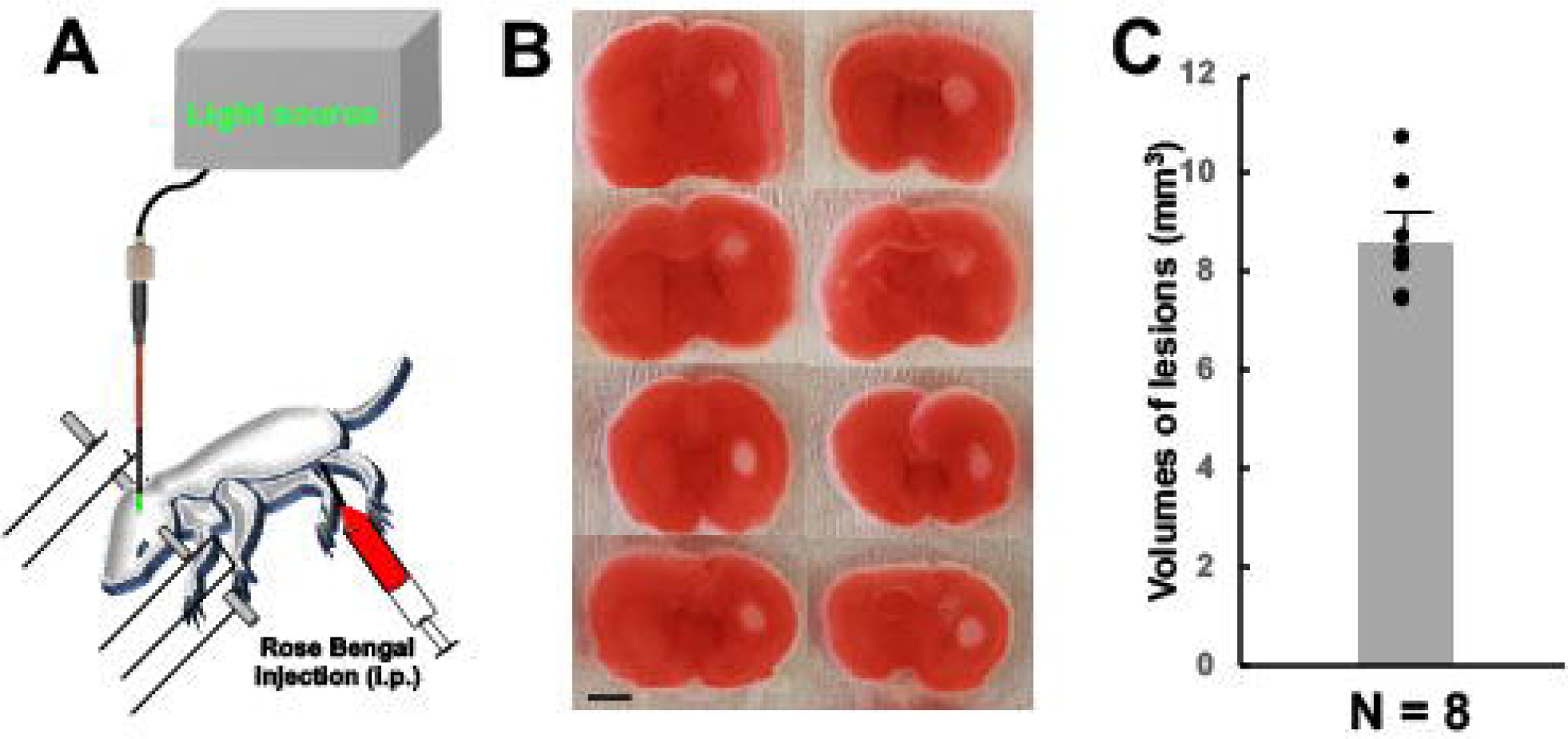
Photothrombotic lesions. **A,** Schematic of photothrombotic method. Lesion was induced by a combination of intraperitoneal injection of Rose Bengal and exposure of green light. **B,** Images of TTC staining at 2 days after lesion. There were 8 rats in total with lesion. The infract lesions were pale white in color. **C,** Volumes of lesions. Bar = 500 μm.

To determine the size and location of the lesions, 2,3,5-Triphenyltetrazolium chloride (TTC) staining was performed, which identifies infarcted brain (white) versus healthy brain (red; Figure 3B). The rat pups were anesthetized with pentobarbital and decapitated 2 days after photothrombotic lesions. The brain was removed and sectioned coronally into five 1-mm slices in a brain matrix. Slices were incubated in 0.1% TTC (Sigma-Aldrich, St. Louis, MO, USA) solution at 37 °C for 20 min and fixed in 10% buffered formalin at 4 °C overnight.

### Electrolytic lesion

The electrolytic lesion was performed in P7 rat pups (N=20; 12 for anatomical analyses and an additional 8 for combined anatomical and behavior experiments) by using a current generating lesion-making device (Model 53500, UGO BASILE S.R.L., VA, Italy). The rat pups were anesthetized using isoflurane as described above. Then, a midline incision was made to expose the midline and Bregma. The target lesion location was the TCT center: – 2.2 mm anterior-posterior (AP), 2.2 mm lateral to midline (ML), and – 4.5 mm deep to the brain surface (dorsal-ventral; DV) according to the stereotaxic coordinates of the P7 rat brain atlas [24]. A burr hole was drilled at this location. Then, a microelectrode of 500 µm in diameter (FHC, Bowdoin, ME, USA) connected with the lesion making device was inserted through the hole to cause electrolytic lesion by using a current of 0.5 mA for 10 s (Figure 4A). In the sham-operated rats (N=6), the microelectrode was inserted, but no current was delivered.

**Figure 4.**
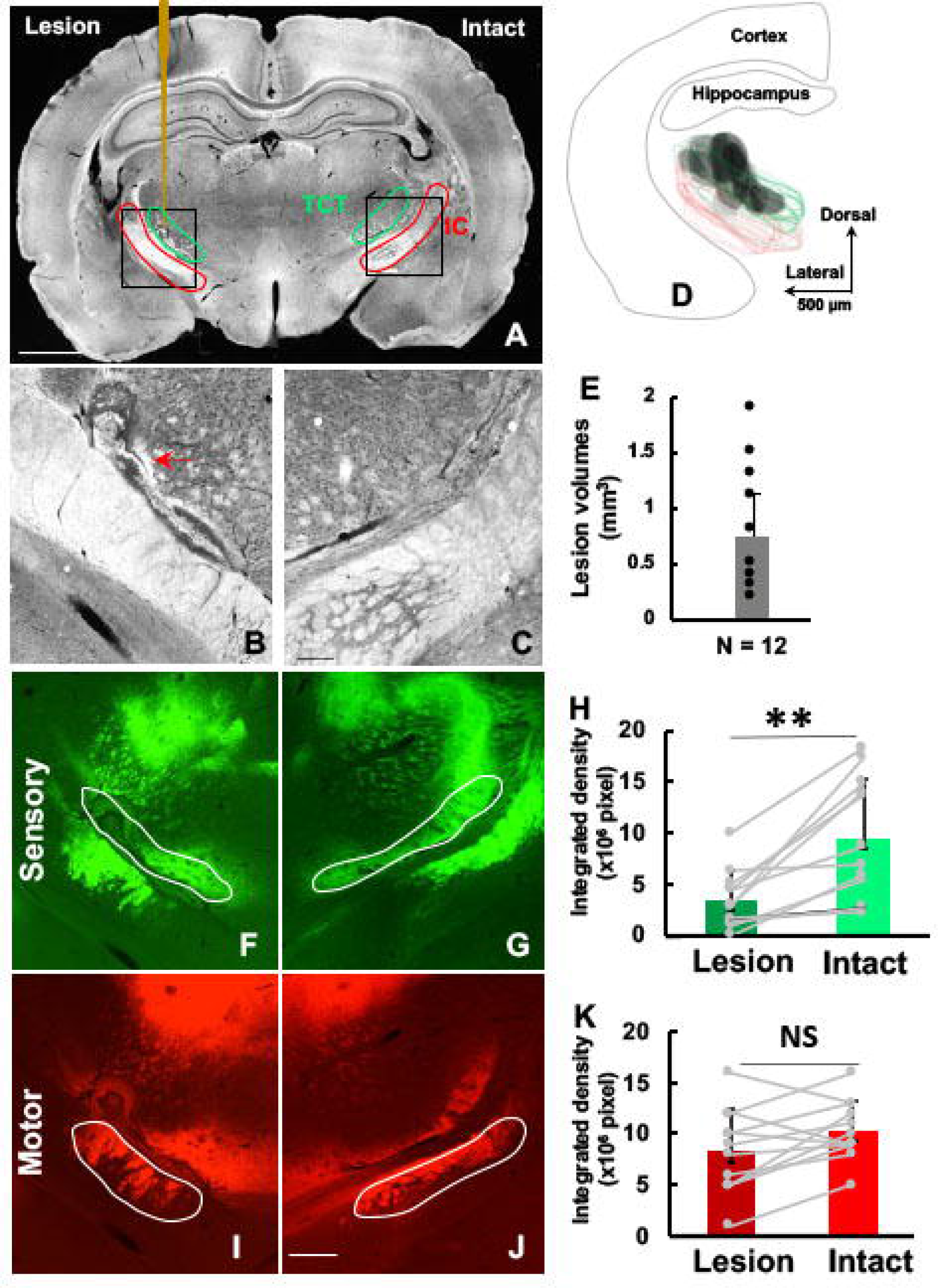
Electrolytic lesions. **A,** A representative image of a coronal section from a rat with the left lesion. Green circles, TCT; Red, IC;, electrolytic probe. **B, C,** High magnification images of the lesion side (red arrow, **B**) and the intact side (**C**)**. D**, Overlap of lesions. Green circles: TCT; Red circles: IC; Grey ones: lesions. **E**, Volumes of lesions. **F, G,** Representative images of viral labeling in TCT of the lesion side (**F**) and the intact side (**G**). **H,** Integrated density of labeling of TCT. **I, J,** Representative images of viral labeling in IC of the lesion side (**I**) and the intact side (**J**). **K,** Integrated density of labeling of CST. Note that there was a significant reduction in TCT (**H**), but not in CST (**K**). **, p < 0.01, NS, no significant. Bar = 500 μm in A, D and E, 125 μm in C and J.

To determine the effects of the lesion on the axon fibers within the TCT, the retrograde adeno-associated virus 2-enhanced green fluorescent protein (AAV2-rGFP, Penn Vector Core, Philadelphia, PA) was injected into the sensorimotor cortex immediately after the lesion induction [25]. Total of 0.2 µl of the retrograde (6 x 10^12^ cfu/µL) was injected into each hemisphere at 0.0 mm AP, 4.0 lateral by using a MicroPump (WPI, Sarasota, FL, USA) at a flow rate of 0.2 µl/min. To determine the effects of the lesion on the axon fibers within the IC, the anterograde tracer AAV8-CAG.CI-rCOMET (AAV8-rCOMET, Penn Vector Core) was injected into the motor cortex immediately after the lesion induction [26]. Total of 0.2 µl of the anterograde tracer (2.4 x 10^12^ cfu/µL) was injected into each hemisphere at 1.0 mm AP, 1.5 mm lateral by using the MicroPump at a flow rate of 0.2 µl/min. Finally, the skin was closed using topical tissue adhesive (WPI), and the pups were warmed on a heating plate until fully awake before being returned to their dams.

Two weeks after the tracer injection, rats were perfused as described above. Brains were removed, postfixed overnight, and then transferred to a solution of 30% sucrose in 0.1 M PBS for 3 days. The region of the brain including the lesion was blocked, frozen with dry ice, and kept at −20 ℃ until sectioning. Coronal sections (40 μm) were cut on a cryostat, and every tenth section was collected from 2.0 to −3.0 mm AP (12 total) for analysis.

Photomicrographs were captured using a Zyla sCMOS camera (Andor-Oxford Instruments, Tubney Woods, Abingdon, UK), mounted to a Nikon ECLIPSE Ni microscope (Nikon Instruments Inc., Melville, NY, USA). Lesions in the left TCT and anatomic structures such as cortex, hippocampus, and ventricles in both hemispheres were traced using NeuroLucida software (MBF Bioscience, Williston, VT) under darkfield at 40x magnification. The viral tracer-labeled areas in the left or right TCT were traced using fluorescence imaging at 100x magnification. Then, the traced structures were reconstructed using NeuroLucida software (MBF). For volumes and coordinates of the lesion, we used NIS Elements AR software (Nikon Instruments Inc.) and the stereotaxic coordinates of the P21 rat brain atlas [24]. In addition, integrated density of the tracer labeled areas in the TCT of both hemispheres were measured.

### Behavioral tests

#### Pasta Handling

From 8 weeks after electrolytic lesion, the rats (N=8, 4 males, 4 females) were videoed while they ate uncooked pieces of pasta, and the number of forepaw adjustments were counted during video replay at ¼ speed. Vermicelli pasta was cut into 7 cm lengths and scores from at least 5 pieces of pasta were averaged for each animal [7, 16, 27]. Testing was performed in an apparatus like the home cage with clear walls, and the rats were habituated to eating in front of the experimenter for at least 3 days prior to initiation of testing.

#### Cylinder exploration test

To assess spontaneous forelimb use, the rats were placed in a clear Plexiglas cylinder (40 cm tall and 20 cm in diameter) with two mirrors placed behind the cylinder to observe forelimb use. The cylinder task was always performed before the pasta handling. The rats were videoed while freely exploring the cylinder for five minutes and were scored based on the number of touches they used for each forelimb to the side of the cylinder while rearing to explore.

### Statistics

Statistical analyses were performed using SPSS (version 22; Chicago, IL). Data points were shown as mean ± SD. The differences in intensity of the tracer labeling in the cortex and in TCT were tested by performing Student t-test. Data for pasta manipulations and cylinder exploration tests were tested by performing repeated measures 1-way ANOVA followed by Bonferroni correction for multiple comparisons (2 behavioral tasks). The significance was set to p < 0.05.

## Results

### Modeling injury to the thalamocortical tract

We developed clinically relevant models of unilateral sensory system injury. The three main components of human injury we sought to incorporate were pathophysiology, location, and selectivity. Since most periventricular lesions leading to hemiplegia are caused by periventricular hemorrhagic infarction, we started with blood injection in the periventricular white matter in the TCT location. All lesions were performed at P7.

### Blood injection

Twenty P7 rats received blood injections under ultrasound guidance (Figure 2). Four of the rats were perfused at 3 days after injection to check the size and position of the hematoma. The other 16 rats were perfused at 12 weeks after blood injection to measure the lesions. In the Nissl-stained coronal sections, we observed hematoma in all of the 4 rats at 3 days after blood injection (Figure 2D). However, none of the rats had lesions at 12 weeks after blood injection (Figure 2E, 2F). In addition, we measured the volumes of both hemispheres to determine changes in hemisphere volume at 12 weeks after blood injection. The was no difference in the volume of the two hemispheres; the left hemisphere (injected side) was 0.967 ± 0.081 cm^3^, and the right side (no injection) was 0.957 ± 0.108 cm^3^ (Figure 2G, p > 0.05). Given the lack of long-term injury with the blood injection model, we pursued alternative approaches to induce persistent lesions.

### Photothrombotic lesions

Photothrombosis with combination of injection of Rose Bengal and exposure of green light was used to induce a lesion in the periventricular area (Figure 3A). In the lesioned hemisphere, the infarct in all of 8 rats were white in color, and the intact tissues were red (Figure 3B). TTC staining also showed that the lesions were in the targeted areas, and the sizes of the lesions were similar in all 8 rats. However, the mean infarct volume was relatively large (Figure 3C, 8.6 ± 1.1 mm^3^) and it caused injury not only to TCT, but also to IC (motor) and other structures. Therefore, we had to forgo this lesion model due to lack of specificity for targeting the sensory tract.

### Electrolytic lesion

Using the optimized electrode, current and duration (see Methods), all 12 rats had lesions largely contained within the center of TCT on the left lesion side (Figure 4A, 4B). Lesions were detected within the TCT in 9 rats, 2 rats had lesions just dorsal to the TCT and near the lateral ventricle, whereas one rat had a lesion that extended into the IC. In the control rats (N=6) that had the electrode position, but no current delivered, we did not detect any damage in the targeted TCT. On the side without electrode penetration, we also did not detect any damage (Figure 4C). To demonstrate the extent of injury, we overlaid the coronal section with the largest lesion from each injured rat; the lesions largely overlapped among these rats (Figure 4D). The average volume of the lesions was 0.75 ± 0.6 mm^3^ (Figure 4E). The average coordinates for the lesion across the rats were AP: 2.6 ± 0.5 mm, ML: 3.4 ± 0.5 mm, and DV: 4.5 ± 0.5 mm, suggesting that lesions were located close to the target.

We used viral tracers to quantify changes of the TCT and CST after lesions in the rats. The retrograde tracer AAV2-rGFP was used to label the TCT as green (Figure 4F and 4G), and the anterograde tracer AAV8-rCOMET to label the CST as red (Figure 4I and 4J). In comparison to the intact right side, more than half of the TCT was lost in the lesioned left side of the rats, only 37.5 ± 12.0% of the TCT was detected in the lesioned left side (Figure 4F and 4G). The mean integrated intensity of the right intact side was 9.4 ± 6.0 x10^6^ and significantly larger than the left injured side 3.4 ± 6.0 x10^6^ (p < 0.01, Figure 4H). However, the tracer intensity of the CST was not significantly different on both sides of the IC (Figure 4I and 4J). The mean integrated intensity of the right intact side was 10.3 ± 2.9 x10^6^, the intensity of the left injured side was 8.3 ± 4.1 x10^6^ (p > 0.05, Figure 4K).

### Behavioral test

After confirming the ability to specifically and accurately target the TCT sensory tracts with electrolytic lesions, we tested the hypothesis that lesions to the sensory circuit will result in impaired forelimb use due to a decrease in sensory input. Adult rats (N=8, 4 Male and 4 Female) that had undergone electrolytic lesions of the TCT at P7 had testing on two behavioral tasks to assess forelimb function. Using the cylinder behavioral task, we found that animals had a significant reduction in the use of their contralesional (impaired) forelimb when compared to the ipsilesional forelimb (Figure 5A, p < 0.01). Interestingly, we did not find a difference in the number of adjustments made by each forelimb in the pasta handling task (Figure 5B), suggesting that the motor deficits after TCT were subtle enough to only affect some behavioral tasks. After performing the behavioral experiments, we visualized the electrolytic lesions and confirmed TCT lesions in 7/8 of these rats (Figure 5C). We found that the electrolytic lesions in the behavior animals specifically targeted the sensory circuits with a reduction of the TCT on the lesion side compared to the intact side (Figure 5D, p < 0.01), while the motor CST tracts were unchanged on the lesion and intact side (Figure 5E). Finally, we found that there was a stronger correlation between the density of the labeled axons in the lesioned side and the percent use of the impaired paws in the cylinder task (r = 0.576, p < 0.05, Figure 5F), indicating that the behavioral deficits in forelimb use was associated with specific injury of the sensory tract.

**Figure 5.**
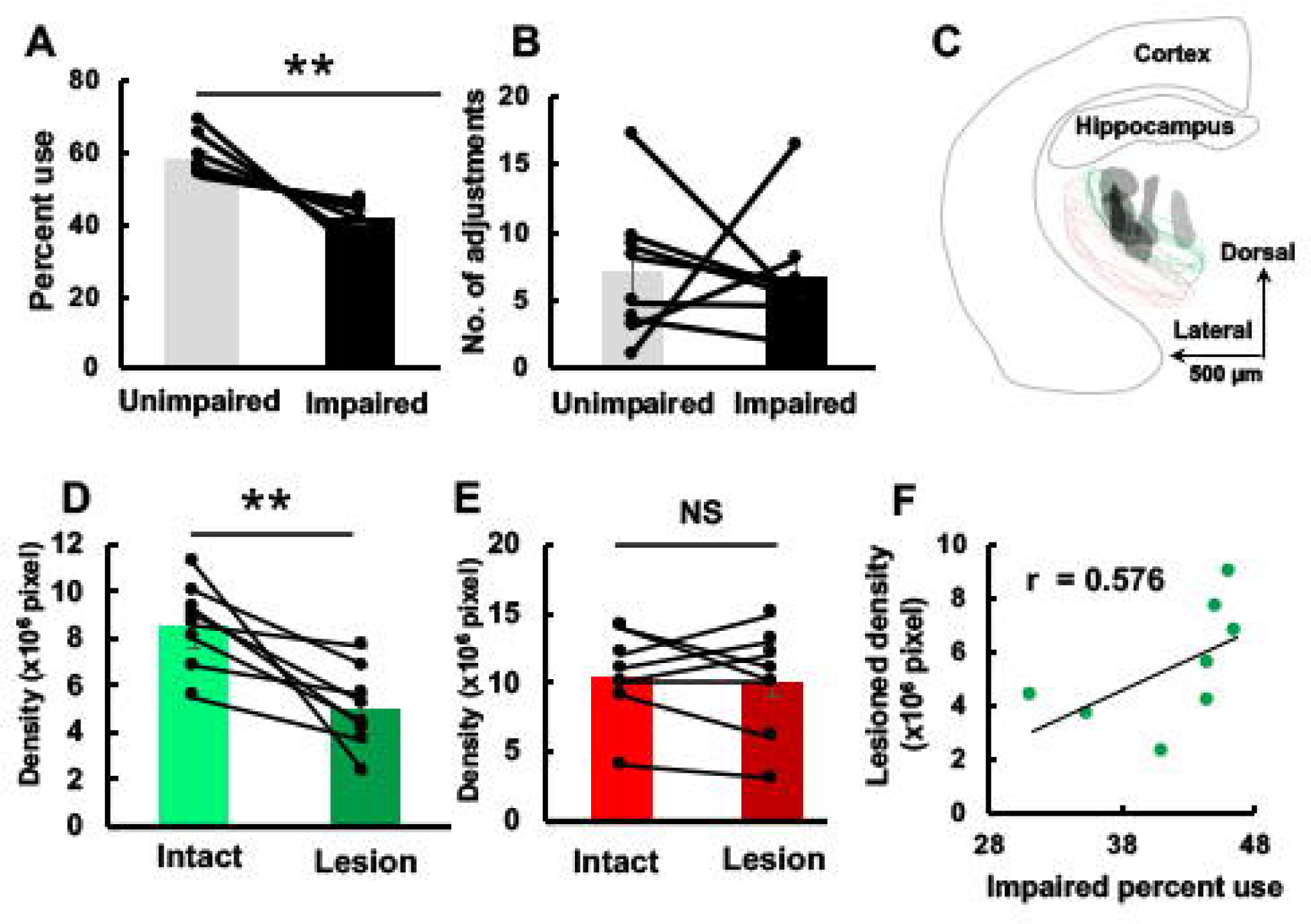
Behavioral forelimb use and anatomical analysis. **A,** Percent forelimb use on the cylinder test. **B,** Forelimb adjustments on the pasta handling task. **C,** Overlap of lesions of behavior rats. **D, E,** Integrated density of labeling of TCT (**D**) and CST (**E**). Note that there was a significant reduction in TCT (**D**), but not in CST (**E**). **F,** Correlation between the density of the labeled axons in the lesioned side and the percent use of the impaired paws in the cylinder task. ** p < 0.01, NS, no significant. Bar = 500 μm.

## Discussion

We aimed to develop a reliable injury model that specifically targeted sensory circuits to investigate the role of sensory transmission on skilled movement. The development of the neonatal stroke model had to overcome specific challenges such as accurate targeting in a neonatal brain, specifically targeting a sensory tract that was in close proximity to the motor tract. Three methods were tested to establish a neonatal sensory injury model: 1) blood injection method, 2) photothrombotic lesion, and 3) electrolytic lesion. The electrolytic model was accurate in its targeting and reproducible between subjects.

The blood injection model can be used to mimic intracerebral hemorrhage [28]. Benefits of the blood injection model include mimicking the blood toxicity effects observed after clinical hemorrhagic strokes [29]. Limitations to the blood injection model include variable lesion size across subjects (Figure 2). Additionally, it has been noted that this method has limited use in long-term studies [30], and we found that the lesion did not remain at 12 weeks.

The photothrombotic method was used to induce ischemic stroke and was useful in producing reproducible lesions to large regions of the brain [30]. This method mimics some mechanisms disrupted in clinical models of stroke, including reactive oxygen species induction, damage to the endothelium, and platelet aggregation [29]. Additionally, this method has the benefit of requiring minimally invasive surgery [30]. Limitations of this method for the current study is the induction of a large lesion that affected both the sensory and motor circuits (Figure 3).

The electrolytic lesion was the most suitable for the current study to investigate the specific role of sensory circuits in motor function after early brain injury. This method allowed for a small precise lesion to TCT while sparing the motor tracts. The electrolytic lesion was also highly reproducible across subjects. Disadvantages of this approach include decreased clinical translational ability given the difference in mechanisms of injury. Additionally, although we were aiming for a small precise lesion, the method may not be appropriate for studies aiming to mimic large cerebral vascular territory injury.

In terms of behavior, we found that electrolytic lesions of TCT also impacted behavior, specifically forelimb preference in the cylinder task. We did not find a difference in forelimb preference in pasta handling suggesting a relatively mild injury, as predicted by the small size of less than a cubic millimeter. Interestingly, we found a correlation between behavioral deficits in the cylinder task and intensity of thalamocortical axon density labeling, providing further evidence that sensory circuits contribute to skilled movement after early brain injury. Overall, the data suggest the TCT contributes to motor behavior and sensory pathways play a role in motor outcomes after early brain injury. Therefore, targeting sensory pathways for repair after injury has the potential to contribute to meaningful movement recovery.

In humans, it was found that a combination of motor and sensory circuit disruptions impairs movement control more than pure motor circuit disruption [10]. In this study, we specially targeted the sensory system and found that targeted lesions to the sensory system impair motor behavior. Interestingly, not all movement is impaired, we found disruptions in forelimb preference in free exploration of the environment but not with pasta manipulation. These results correlate with human findings that show specific domains of motor performance are impaired depending on the type of brain injury and therapeutic interventions[31]. The findings suggest that the TCT plays a role in motor behavior, specifically forelimb preference in free exploration, or that exploration is a more sensitive measure of these partial lesions.

Future experiments will be aimed at specifically manipulating TCT with neuromodulation interventions to demonstrate the sufficiency of sensory repair for movement recovery. For example, an advantage and limitation of the current study was that only a small proportion of the TCT was disrupted with the electrolytic lesion technique. Therefore, a more complete lesion of the tract may result in larger behavioral deficits. Additionally, given that we did see behavioral consequences with the small lesion, the spared region of the tract could be stimulated to investigate the effects of sensory tract activation on movement recovery.

## Study approval statement

The study was done in accordance with the National Institutes of Health guidelines recommendations and approved by the Institutional Animal Care and Use Committee of Columbia University. The approval number is AC-AABP2552.

## Conflict of interest statement

The authors have no conflicts of interest to declare.

## Funding sources

Weinberg Family Cerebral Palsy Center, Department of Neurology and R01NS115470. The funder had no role in the design, data collection, data analysis, and reporting of this study.

## Author contributions

JBC, the study design, and manuscript conception, planning, writing and decision to publish. MC, the study performance, data analysis, and manuscript writing. TCW, the study design and performance, data analysis, and manuscript writing.

## Data Availability statement

Summary data are presented in the article, and source data can be requested from the corresponding author.

## Acknowledgements

We thank A. Ramamurthy for assistance in scoring the pasta handling behavioral data. We thank members of the Carmel lab for their helpful suggestions.

## Notes

### Competing Interest Statement

The authors have declared no competing interest.

